# Model-based and model-free analyses of the neural correlates of tongue movements

**DOI:** 10.1101/812263

**Authors:** Peter Sörös, Sarah Schäfer, Karsten Witt

## Abstract

The tongue performs movements in all directions to subserve its diverse functions in chewing, swallowing, and speech production. The aims of the present study were twofold: using task-based functional MRI in a group of 17 healthy young participants, we studied (1) potential differences in the cerebral control of frontal, horizontal, and vertical tongue movements and (2) potential inter-individual differences in tongue motor control. To investigate differences between different tongue movements, we performed voxel-wise multiple linear regressions. To investigate inter-individual differences, we applied a novel approach, *spatio-temporal filtering of independent components.* For this approach, individual functional data sets were decomposed into spatially independent components and corresponding time courses using ICA. A temporal filter (correlation with the expected brain response) was used to identify independent components time-locked to the tongue motor task. A spatial filter (cross-correlation with established neurofunctional systems) was used to identify brain activity not time-locked to the task. Our results confirm the importance of an extended bilateral cortical and subcortical network for the control of tongue movements, including the lateral primary sensorimotor cortex, supplementary motor cortex, anterior cingulate gyrus, insula, basal ganglia, thalamus, and cerebellum. Frontal tongue movements, highly overlearned movements related to speech production, showed less activity in parts of the frontal and parietal lobes compared to horizontal and vertical movements and greater activity in parts of the left frontal and temporal lobes compared to vertical movements. The investigation of inter-individual differences revealed a component representing the tongue primary sensorimotor cortex time-locked to the task in all participants. Using the spatial filter, we found the default mode network in 16 of 17 participants, the left fronto-parietal network in 16, the right fronto-parietal network in 8, and the executive control network in 4 participants. Further detailed analyses of speech-related tongue movements are warranted to increase our understanding of tongue motor control as a crucial part of articulation. Spatio-temporal filtering of independent components appears to be a powerful approach to study inter-individual differences in task-based functional MRI. This approach may be particularly useful for the assessment of individual patients and may be related to individual clinical, behavioral, and genetic information.

## 1 INTRODUCTION

The human tongue is a unique muscular and sensory organ with critical roles in several motor tasks, such as chewing, swallowing, respiration, and speech (Hiiemae and Palmer, 2003; Sawczuk and Mosier, 2001), in addition to its somatosensory (Pardo et al., 1997; Sakamoto et al., 2010) and gustatory functions (Hummel et al., 2010; Kobayakawa et al., 2005).

To subserve its distinct motor tasks, the tongue contains intrinsic and extrinsic muscle fibers (Abd-El-Malek, 1939; Schumacher, 1927), which are extensively interwoven (Gaige et al., 2007). Intrinsic fibers originate and insert within the tongue itself, while extrinsic fibers are attached to bony structures such as the mandible, hyoid bone, or styloid process (Sanders and Mu, 2013). This complex biomechanical architecture is the basis for the tongue’s ability to move and alter its shape in all three dimensions. Moreover, adult human tongues, compared to the tongues of other mammals, are characterized by a higher proportion of slow-twitch (type I) muscle fibers, which are associated with fine motor control (Sanders et al., 2013). Intrinsic and extrinsic tongue muscles are innervated by the lateral and medial divisions of the hypoglossal nerve (cranial nerve XII), with different components of the musculature being supplied by different hypoglossal branches (Mu and Sanders, 2010) and controlled by distinct hypoglossal subnuclei (McClung and Goldberg, 2002).

The cortical and subcortical control of tongue movements has been studied thoroughly in animals and humans using various invasive and non-invasive techniques. These electrophysiologic, neuroimaging and lesion studies suggest that voluntary (e.g. speech-related) and semi-automatic (e.g. swallowing-related) tongue movements (Martin et al., 1997) are controlled by the lateral primary sensorimotor cortex (Takai et al., 2010), supplementary motor area, basal ganglia, and cerebellum (Corfield et al., 1999; Martin et al., 2004; Shinagawa et al., 2003; Watanabe et al., 2004).

Using functional magnetic resonance imaging (fMRI), researchers investigated (1) isolated voluntary tongue movements, such as tongue protrusion (Arima et al., 2011), horizontal tongue movements (Riecker et al., 2000), and tongue elevation (Martin et al., 2004), and (2) tongue movements as part of speaking (Riecker et al., 2005; Sörös et al., 2006), singing (Jungblut et al., 2012; Ozdemir et al., 2006), and swallowing (Sörös et al., 2009; Lowell et al., 2012). A detailed comparison of the neural correlates of different tongue movements in all three directions has not been performed yet (but see the study by Watanabe et al. (2004), comparing tongue protrusions in different directions with tongue retraction). Moreover, almost all fMRI studies on tongue movements present only group analyses (one notable exception is the study by Martin et al. (2004), Table 5, presenting individual brain activation in all studied participants). Finally, almost all fMRI studies on tongue motor control have been performed on older scanner hardware and with relatively small sample sizes.

The first aim of the present study was to identify and compare neural activity associated with three different tongue movements: frontal tongue movements (protrusion), horizontal tongue movements, and vertical tongue movements (elevation). These tongue movements are used in different tongue motor tasks. Frontal tongue movements are almost exclusively used in speech production and singing. Horizontal tongue movements are used during chewing to position the food in the oral cavity and to form the bolus in preparation for swallowing. Vertical tongue movements, finally, are used in both speech production and the oral phase of swallowing.

The second aim of this study was to investigate inter-individual similarities and differences in tongue movement-related neural activity. The number of swallows (Rudney et al., 1995) and of words produced per day (Mehl et al., 2007) varies considerably between individuals. To investigate individual brain activity, we performed an independent component analysis of individual task-based fMRI data sets. With a novel approach, *spatio-temporal filtering of independent components*, we identified individual components whose time courses were highly correlated with the expected brain response and whose spatial patterns were highly correlated with one of the established neurofunctional networks described by Smith et al. (2009).

## 2 METHODS

### 2.1 Participants

Twenty young healthy individuals have been investigated for the present study. Three participants were excluded from the data analysis (2 because of excessive head motion, 1 because of severe artifacts in the first-level analysis). For the final data analysis, the MRI data of 17 participants (9 women, 8 men) have been investigated. Mean ± age standard deviation (SD) was 25. ± 3.3 years (minimum: 20, maximum: 34 years). All participants met the following criteria: (1) no history of neurological disorders (such as dementia, movement disorder, stroke, epilepsy, multiple sclerosis, traumatic brain injury, migraine), psychiatric disorders (such as schizophrenia or major depression), or cancer, (2) no impaired kidney or liver function, (3) no use of psychotropic medication (in particular, antidepressants, antipsychotics, benzodiazepines, and opioids), (4) no substance abuse, (5) no excessive head motion (< 1 mm relative mean displacement and < 3 mm absolute mean displacement) during fMRI. Handedness was determined with the Edinburgh Handedness Inventory - Short Form (Veale, 2014). Right handedness was present in 13 individuals (handedness scores: 62.5 - 100), mixed handedness was found in 4 individuals (handedness scores: 33.3 - 50). Seven participants have had an MRI scan before, 10 participants have never been within an MRI scanner. All participants gave written informed consent for participation in the study. A compensation of 10 € per hour was provided. The study was approved by the Medical Research Ethics Board, University of Oldenburg, Germany.

### 2.2 Experimental paradigm and tongue movements

The data for the present study were collected as part of a larger project on oral and speech-language functions. **Table 1** summarizes the order in which the four different tasks were performed. The total MRI measurement time was approximately 45 min.

**Table 1.** Structural and functional sequences used for the entire project (scan 4: tongue movements).

| No. | Sequence | Time of acquisition |
| --- | --- | --- |
| 1 | T1-weighted MPRAGE | 6:16 min |
| 2 | T2*-weighted (syllable production) | 9:16 min |
| 3 | T2*-weighted (tongue twister) | 8:31 min |
| 4 | T2*-weighted (tongue movements) | 9:21 min |
| 5 | T2*-weighted (sentence production) | 9:14 min |

For the investigation of the neural correlates of tongue movements, participants were visually cued to perform one of three different repetitive tongue movements (**Figure 1**): (1) frontal tongue movements (also known as tongue protrusion); participants were instructed to push the tip of the tongue against the surface of the maxillary incisors and then retract the tongue to the rest position, similar to the English /th/ sound, (2) horizontal tongue movements; participants were instructed to move the tongue against the right and left mandibular premolars, and (3) vertical tongue movements; participants were instructed to elevate the tongue and to press it against the hard palate (the roof of the mouth), similar to the beginning of the oral phase of swallowing (Dodds, 1989). Before the fMRI measurement, all participants underwent a short training session outside the scanner to familiarize themselves with the different tongue movements and the visual cues. For all conditions, participants performed rhythmic and self-paced movements of approximately 1 Hz. Behavioral performance during fMRI was not recorded.

**Figure 1.**
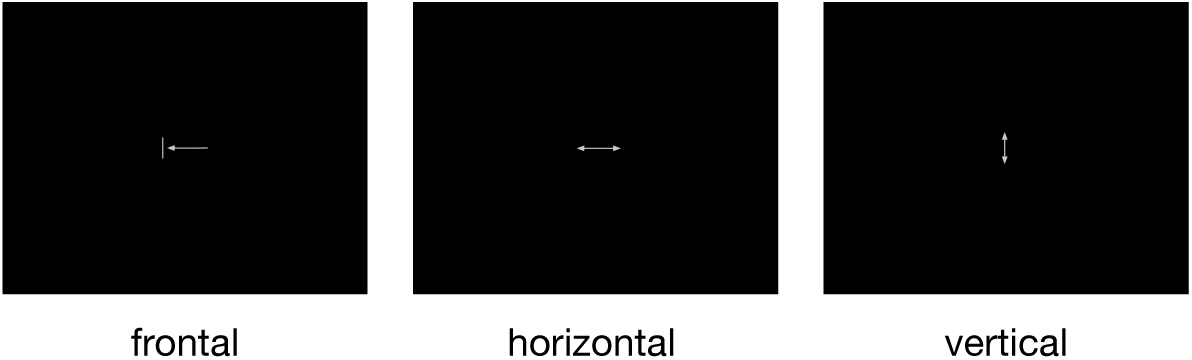
The three pictograms used to cue frontal, horizontal, and vertical tongue movements. All symbols were simple and small to minimize eye movements and visual processing. During rest periods, a fixation cross was shown.

Visual cues were presented by the MATLAB toolbox Cogent Graphics (developed by John Romaya, Laboratory of Neurobiology, Wellcome Department of Imaging Neuroscience, London, UK)^1^ on a PC and projected through an LCD projector onto a screen mounted within the scanner bore behind the head coil. Participants were able to see the cues via a mirror attached to the head coil. Visual cues were shown in blocks of 15 s duration with a 15 s rest condition (during which a fixation cross was presented) after every third tongue movement block. The remaining tongue movement blocks were separated by a shorter rest condition of 3 s. **Figure 2** visualizes the experimental paradigm.

**Figure 2.**
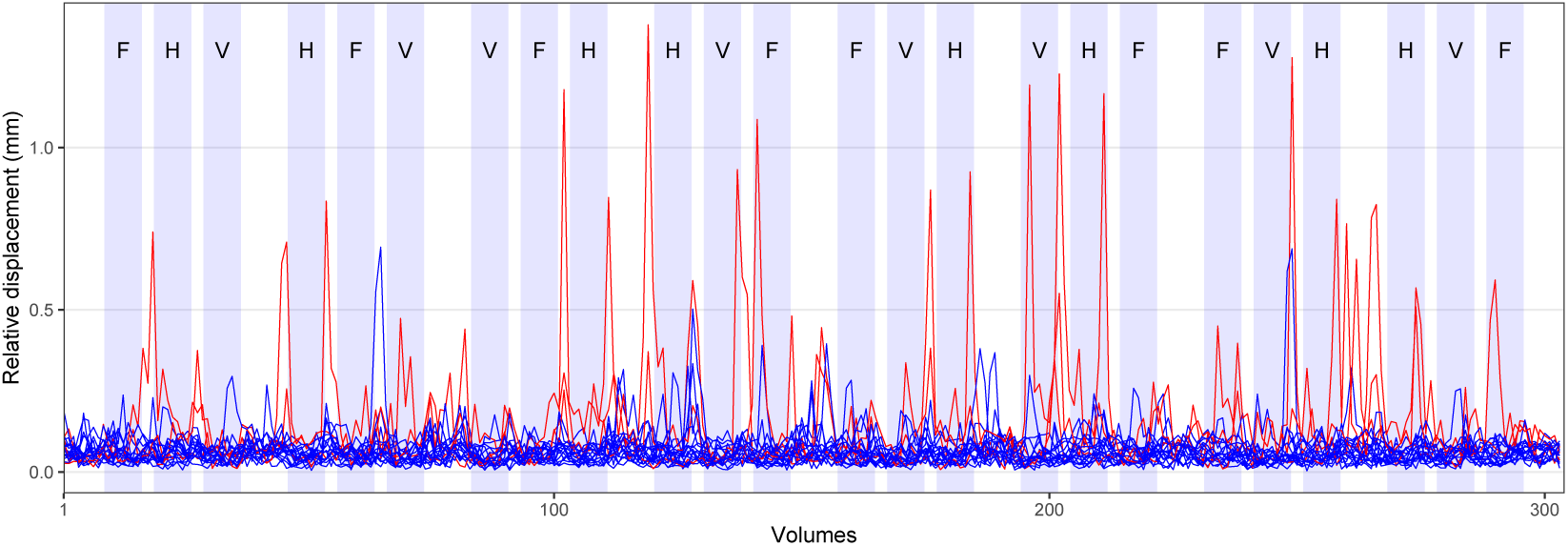
Experimental paradigm and head motion. Blocks of tongue movement are shown in light blue (F: frontal, H: horizontal, and V: vertical movements; duration 15 s each). The mean relative displacement during all fMRI measurements is displayed with blue (17 included participants) and red lines (3 excluded participants).

### 2.3 MRI data acquisition

MR images of the entire brain were acquired at 3 Tesla on a Siemens MAGNETOM Prisma whole-body scanner (Siemens, Erlangen, Germany) with the XR gradient system (gradient strength: 80 mT/m, gradient rise time: 200 T/m/s on all three gradient axes simultaneously) and a 64-channel head/neck receive-array coil. This coil enhances the signal-to-noise ratio of the peripheral image, primarily corresponding to the cortex of the human brain. The scanner is located at the Neuroimaging Unit, School of Medicine and Health Sciences, University of Oldenburg, Germany.^2^

For structural brain imaging, Siemens’ 3-dimensional T1-weighted MPRAGE sequence (Brant-Zawadzki et al., 1992) was used (TR: 2000 ms, TE: 2.07 ms, flip angle 9°, isotropic voxel size: 0.75 × 0.75 × 0.75 mm^3^, 224 axial slices, time of acquisition: 6:16 min). For functional imaging, Siemens’ ep2d bold T2*-weighted gradient-echo echo-planar sequence was used (TR: 1800 ms, TE: 30 ms, flip angle 75°, isotropic voxel size: 3 × 3 × 3 mm^3^, 33 slices, time of acquisition: 9:21 min). Structural and functional measurements used in-plane acceleration (GRAPPA) with a PAD factor of 2 (Lindholm et al., 2009). Before every functional sequence, extended 3-dimensional B0 shimming and true-form B1 shimming was applied. Siemens’ pre-scan normalization filter was deactivated during functional sequences.

### 2.4 MRI data analysis

#### 2.4.1 Preprocessing of structural images

Preprocessing of T1-weighted images was done with the *antsBrainExtraction.sh* script, part of Advanced Normalization Tools (ANTs, version 2.1)^3^ (Tustison et al., 2014). This script performs (1) bias field correction to minimize the effects of magnetic field inhomogeneity using the N4 algorithm (Tustison et al., 2010) and (2) brain extraction using a hybrid segmentation/template-based strategy (Tustison et al., 2014). The script was used together with brain templates derived from the OASIS-1 study.^4^

#### 2.4.2 Preprocessing of functional images

Preprocessing of fMRI data was carried out using FEAT (version 6.00), part of FMRIB’s Software Library (FSL)^5^ (Jenkinson et al., 2012; Smith et al., 2004; Woolrich et al., 2009). Preprocessing included removal of the first 4 recorded volumes to allow for signal equilibration in addition to the 2 dummy volumes measured, but not recorded, as part of the ep2d_bold sequence (304 volumes were retained). Standard head motion correction was performed by volume-realignment to the middle volume using MCFLIRT (Jenkinson et al., 2002). Mean relative displacement (the distance between one volume and the following volume) of all participants is shown in **Figure 2**. Brain extraction of functional images was done with FSL’s brain extraction tool, BET (Smith, 2002). Spatial smoothing with a Gaussian kernel of 5 mm full width at half maximum (FWHM) and grand-mean intensity normalization of the entire dataset by a single multiplicative factor were also performed.

After completion of standard data preprocessing, but without temporal filtering, ICA-based automatic removal of motion artifacts (FSL’s ICA-AROMA version 0.3 beta)^6^ was used to identify and remove motion-related ICA components from fMRI data (Pruim et al., 2015). Here, the non-aggressive option was used, performing a partial component regression. First, ICA-AROMA carries out probabilistic ICA of individual subjects’ MRI data using FSL’s MELODIC (Beckmann and Smith, 2004). Second, ICA-AROMA employs four theoretically motivated temporal and spatial features to select motion-related components from MELODIC’s output. Finally, it removes these components from the initial data set through an ordinary least squares regression using FSL’s *fsl_regfilt* command (Pruim et al., 2015). Decomposition of individual datasets created between 47 and 68 independent components (mean: 60 components). To determine the optimal number of components for every data set, MELODIC uses Bayesian principal component analysis (Beckmann and Smith, 2004). Of these components, between 18 and 39 (mean: 30) components were identified as noise and regressed out. At least for the preprocessing of resting-state fMRI data, ICA-AROMA compares favorably to other methods of head motion correction (Parkes et al., 2018).

Following ICA-AROMA, data were high-pass filtered (Gaussian-weighted least-squares straight-line fitting, sigma = 45 s). Registration of functional to high-resolution structural images was carried out using FLIRT (Jenkinson et al., 2002). Registration from high-resolution structural to Montreal Neurological Institute (MNI152) standard space was further refined using 12-parameter affine transformation and non-linear registration with a warp resolution of 10 mm in FNIRT.^7^

#### 2.4.3 Model-based individual and group fMRI analysis

For first-level model-based analysis, functional data sets were analyzed with a general linear model-based time-series analysis using voxel-wise multiple linear regressions (Friston et al., 1995; Monti, 2011) as implemented in FEAT. The time courses of the three movement conditions were convolved with a gamma hemodynamic response function (using the standard settings: phase: 0 s, standard deviation: 3 s, mean lag: 6 s) and served as regressors of interest. The temporal derivative of each primary regressor was included as a regressor of no interest to improve the model fit when the timing was not exactly correct (e.g., if tongue movements were started or stopped with a slight delay). Regressors of interest (experimental conditions) and regressors of no interest (temporal derivatives) formed the design matrix used for voxel-wise multiple linear regressions. To remove temporal autocorrelations, time-series pre-whitening was used (Woolrich et al., 2001). After generating parameter estimates (PEs) for every primary regressor and every participant, the following contrasts of parameter estimates (COPEs) were calculated: (1) frontal > rest, (2) horizontal > rest, (3) vertical > rest, (4) horizontal > frontal, (5) vertical > frontal, (6) frontal > vertical, (7) horizontal > vertical, (8) frontal > horizontal, and (9) vertical > horizontal.

For higher-level analysis, mixed-effects group analysis maps were generated by FLAME (stage 1 and 2) for all contrasts. FLAME uses a fully Bayesian inference technique in a two-stage process: a fast approach using maximum a posteriori estimates and a slower, more accurate approach using Markov Chain Monte Carlo methods (Woolrich et al., 2004). Z statistic images were thresholded non-parametrically using a cluster-forming threshold of Z > 3.1 and a (corrected) cluster significance threshold of p < 0.01 assuming a Gaussian random field for the Z-statistics. No additional correction for multiple contrasts was performed. Local maxima were identified within the Z statistic images using FSL’s *cluster* command (maximum number of local maxima: 100, minimum distance between local maxima: 20 mm). The anatomical location of each local maximum was determined with FSL’s *atlasquery* command and the following probabilistic atlases:^8^ (1) Harvard-Oxford cortical structural atlas (48 cortical areas), (2) Harvard-Oxford subcortical structural atlas (21 subcortical areas), and (3) Probabilistic cerebellar atlas (28 regions) (Diedrichsen et al., 2009).

#### 2.4.4 Model-free individual fMRI analysis

A single-session probabilistic ICA was conducted to decompose every preprocessed individual fMRI data set into 20 independent spatial components and corresponding time courses using MELODIC (version 3.14). The data sets fed into ICA were the preprocessed data sets used for first-level model-based analysis (i.e., after denoising with ICA-AROMA). Of note, these data sets contained brain activity associated with all three tongue movements. A low-dimensional decomposition into 20 components was chosen for later comparison with a well-established set of published networks. MELODIC performs linear decomposition (Comon, 1994) of the original fMRI signal using the FastICA technique (Hyvärinen, 1999) and variance-normalization of the associated timecourses. Spatial maps were thresholded using a Gaussian mixture model approach at a posterior probability level of p < 0.5 (Beckmann and Smith, 2004).

#### 2.4.5 Spatio-temporal filtering of independent components

To identify spatial components whose activity was time-locked to tongue movements, the time course of the expected brain response was correlated with all ICA time courses of all participants. Similarly, Kokkonen et al. (2009) analyzed task-based fMRI activity associated with left and right finger tapping with ICA and correlated ICA time courses with motor task timing. In our study, a significant (p < 0.05) Pearson’s correlation coefficient of r > 0.4 or r < −0.4 between time courses indicated temporal significance.

To identify relevant spatial components not time-locked to the tongue motor tasks, the spatial maps obtained for every participant were cross-correlated with 10 established neural networks^9^ (Smith et al., 2009). A significant Pearson’s correlation coefficient of r > 0.4 between components indicated spatial significance. The cross-correlation of study-specific spatial components with established networks (e.g. developed by Smith et al. (2009) or Yeo et al. (2011)) has been performed previously for the analysis of resting-state fMRI data (e.g., by Reineberg et al. (2015) and Sörös et al. (2019)).

## 3 RESULTS

### 3.1 Head motion

Across all included participants, the average mean absolute displacement of the head (relative to the middle volume) was 0.7 mm (SD: 0.4 mm, min: 0.2 mm, max: 1.9 mm). The average mean relative displacement (compared to the following volume) was 0.3 mm (SD: 0.2 mm, min: 0.1 mm, max: 0.7 mm). The time course of the mean relative displacement of all participants is shown in **Figure 2** (17 included participants in blue, 3 excluded participants in red).

### 3.2 Model-based group fMRI analysis

Compared to rest, the three tongue movements were associated with similar bilateral brain activity (**Figure 3A**). Major regions of cortical activation were the lateral pre- and postcentral gyrus, supplementary motor cortex, anterior cingulate gyrus, and the frontal cortex of both hemispheres. Activation was also found in the bilateral insulae, basal ganglia, thalamus, amygdala, and cerebellum. **Table 2** lists the coordinates in MNI space and the corresponding Z value of significant local maxima of brain activation for the contrast frontal tongue movements > rest.

**Table 2.**
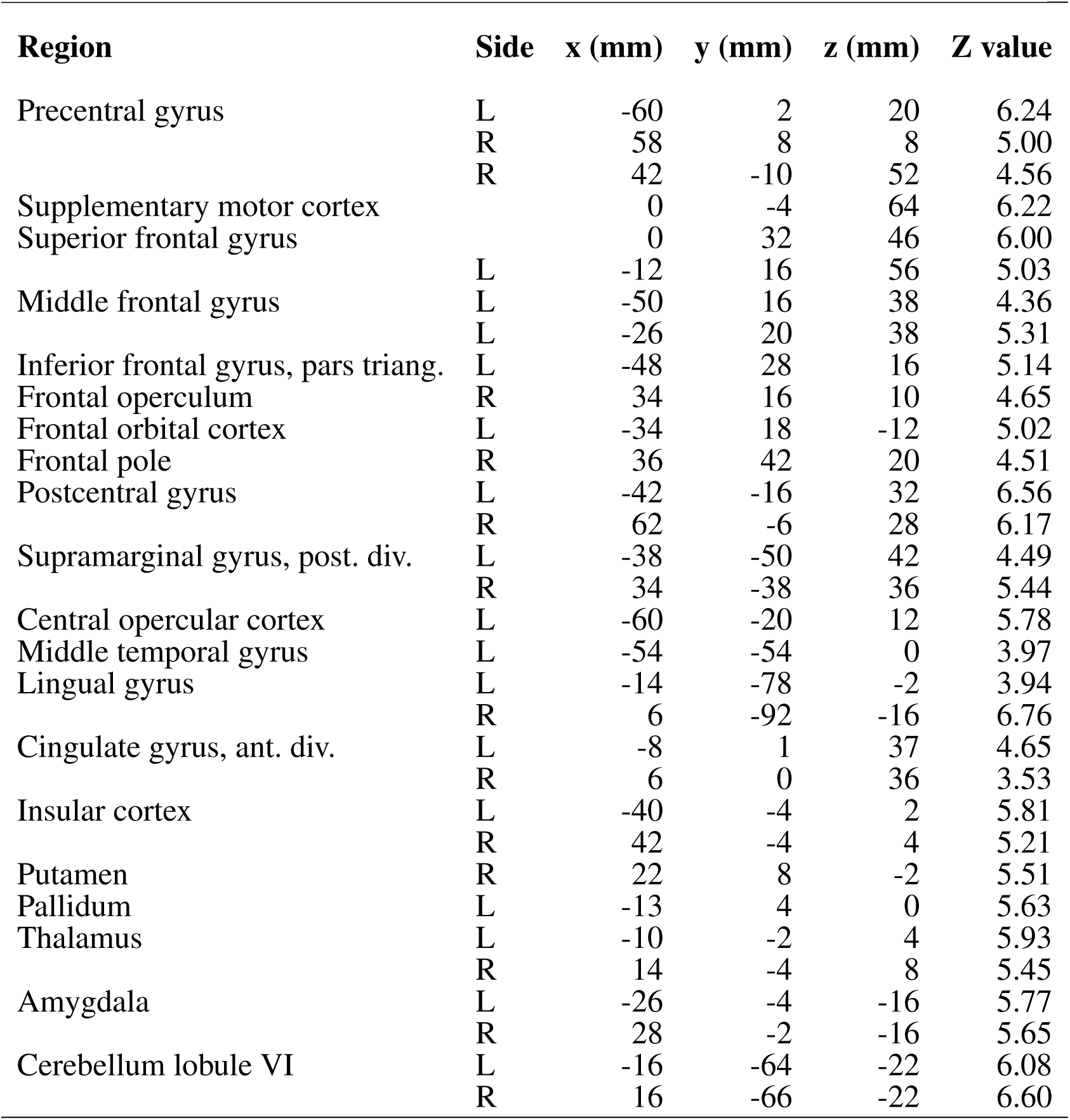
Local maxima in brain activation: stereotaxic coordinates in MNI space, Z values, and corresponding brain regions for the contrast frontal tongue movement > rest.

**Figure 3.**
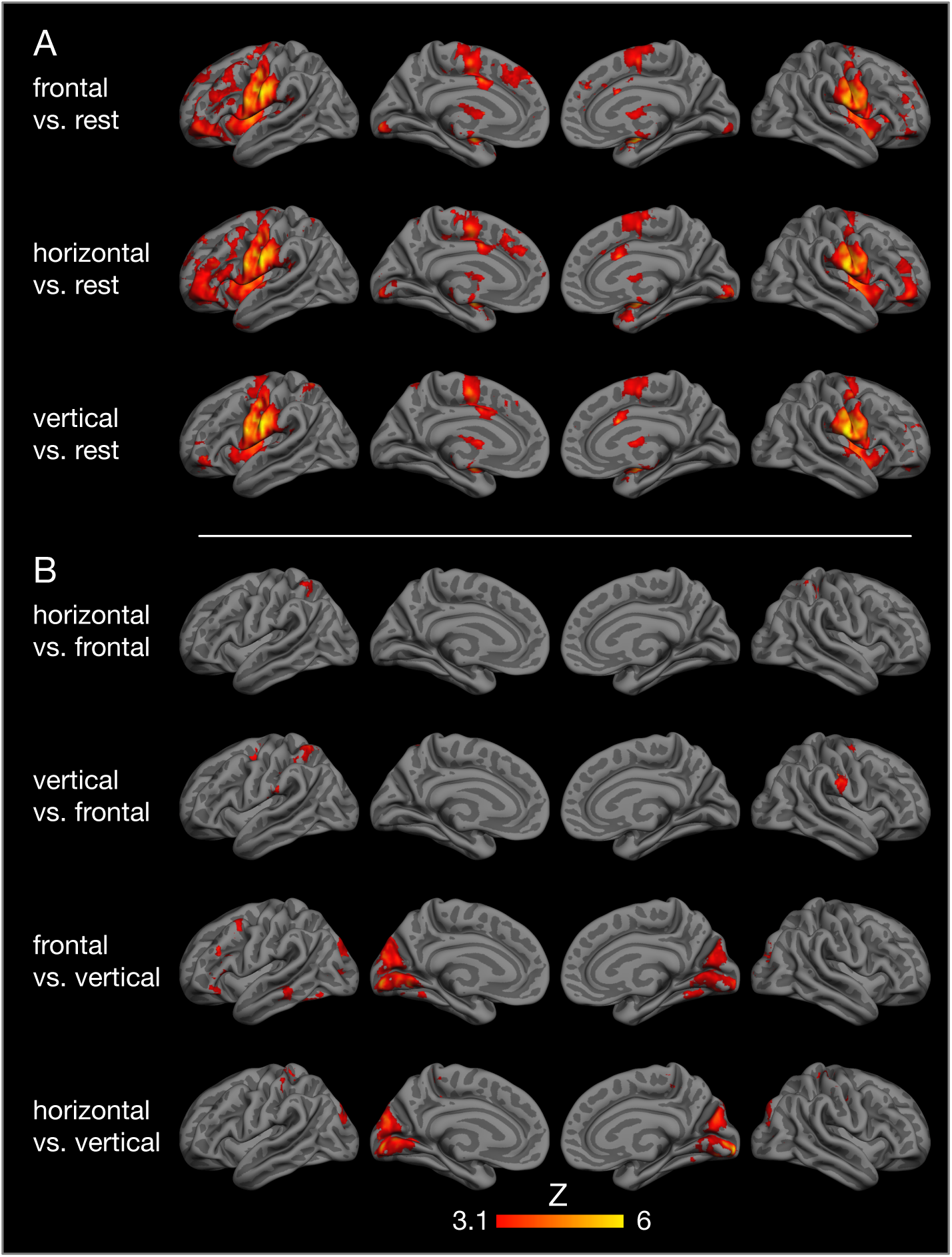
Results of model-based group analysis. Brain activation associated with tongue movements averaged across 17 participants after cluster-based thresholding (Z > 3.1, p < 0.01) is shown in red-yellow. Activated areas were projected onto the semi-inflated pial surface of the *fsaverage* brain, reconstructed by FreeSurfer (Fischl, 2012). (**A**) Tongue movements > rest, (**B**) contrasts between tongue movements.

Contrasts between different tongue movements resulted in significant activation in several cortical areas (**Figure 3B, Table 3**). Horizontal tongue movements were associated with greater activation in the bilateral superior parietal lobule (vs. frontal movements) and in the bilateral pre- and postcentral gyri as well as in the left inferior frontal gyrus (vs. vertical movements). The contrast vertical > frontal movements demonstrated greater activation in the bilateral precentral and the right postcentral gyri, as well as the left supramarginal gyrus and the left superior parietal lobule. The reverse contrast, frontal > vertical movements, showed greater activation in the left frontal and temporal lobes. The contrasts frontal > horizontal and vertical > horizontal did not result in significant activation.

**Table 3.** Local maxima in brain activation: stereotaxic coordinates in MNI space, Z values, and corresponding brain regions for the contrasts horizontal > frontal, vertical > frontal, frontal > vertical, and horizontal > vertical tongue movements.

| Region | Side | x (mm) | y (mm) | z (mm) | Z value |
| --- | --- | --- | --- | --- | --- |
| **Horizontal > frontal** |  |  |  |  |  |
| Superior parietal lobule | L | -32 | -50 | 58 | 4.75 |
|  | R | 26 | -48 | 66 | 4.98 |
| **Vertical > frontal** |  |  |  |  |  |
| Precentral gyrus | L | -26 | -12 | 62 | 5.29 |
|  | R | 32 | -6 | 66 | 4.59 |
| Postcentral gyrus | R | 52 | -20 | 34 | 6.70 |
|  | R | 32 | -34 | 36 | 4.67 |
| Central opercular cortex | L | -60 | -20 | 16 | 4.72 |
| Supramarginal gyrus, ant. div. | L | -44 | -34 | 38 | 5.03 |
| Superior parietal lobule | L | -34 | -48 | 52 | 4.69 |
| **Frontal > vertical** |  |  |  |  |  |
| Middle frontal gyrus | L | -46 | 10 | 46 | 4.56 |
| Inferior frontal gyrus, pars triang. | L | -46 | 24 | 12 | 4.95 |
| Frontal orbital cortex | L | -42 | 30 | -16 | 4.29 |
| Middle temporal gyrus, post. div. | L | -60 | -36 | -10 | 4.33 |
| Inferior temporal gyrus, temp-occ. | L | -42 | -50 | -16 | 4.47 |
| Cuneal cortex | R | 2 | -78 | 22 | 5.56 |
| Lateral occipital cortex | L | -20 | -86 | 20 | 4.79 |
|  | R | 28 | -82 | 12 | 4.85 |
| Intracalcarine cortex | R | 8 | -64 | 8 | 4.45 |
| Lingual gyrus | L | -10 | -78 | 0 | 5.35 |
| Occipital pole |  | 0 | -96 | -4 | 6.33 |
|  | R | 24 | -96 | -8 | 3.87 |
| Occipital fusiform gyrus | L | -32 | -72 | -12 | 4.29 |
|  | R | 26 | -66 | -6 | 4.97 |
| **Horizontal > vertical** |  |  |  |  |  |
| Precentral gyrus | L | -22 | -26 | 58 | 4.74 |
|  | L | -40 | -16 | 46 | 4.60 |
|  | R | 22 | -28 | 68 | 4.97 |
| Inferior frontal gyrus, pars triang. | L | -54 | 28 | -10 | 4.53 |
| Postcentral gyrus | L | -2 | -38 | 58 | 4.83 |
|  | R | 38 | -26 | 50 | 4.67 |
| Cingulate gyrus, post. div. | L | -2 | -20 | 44 | 4.23 |
| Cuneal cortex | L | -6 | -88 | 22 | 5.05 |
|  | R | 10 | -70 | 20 | 5.17 |
| Occipital pole | L | -10 | -90 | -2 | 5.65 |
|  | R | 16 | -98 | -6 | 8.61 |
| Fusiform cortex | R | 26 | -60 | -16 | 5.16 |

### 3.3 Model-free individual fMRI analysis

The individual brain activity of all 17 participants is presented in **Figure 4**. Temporally filtered (time-locked) independent components are shown on the left (**Figure 4A**), spatially filtered components on the right (**Figure 4B**). In all participants, the fMRI data set contained one bilateral sensorimotor component (**Figure 4**, first column) positively correlated with the expected brain activity (graph in the first row), representing the tongue primary sensorimotor cortex. In four participants, another frontal or parietal component (**Figure 4**, second column) was positively correlated with the expected brain activity. In seven participants, a bilateral occipital component (**Figure 4**, third column) was negatively correlated with the expected brain activity.

**Figure 4.**
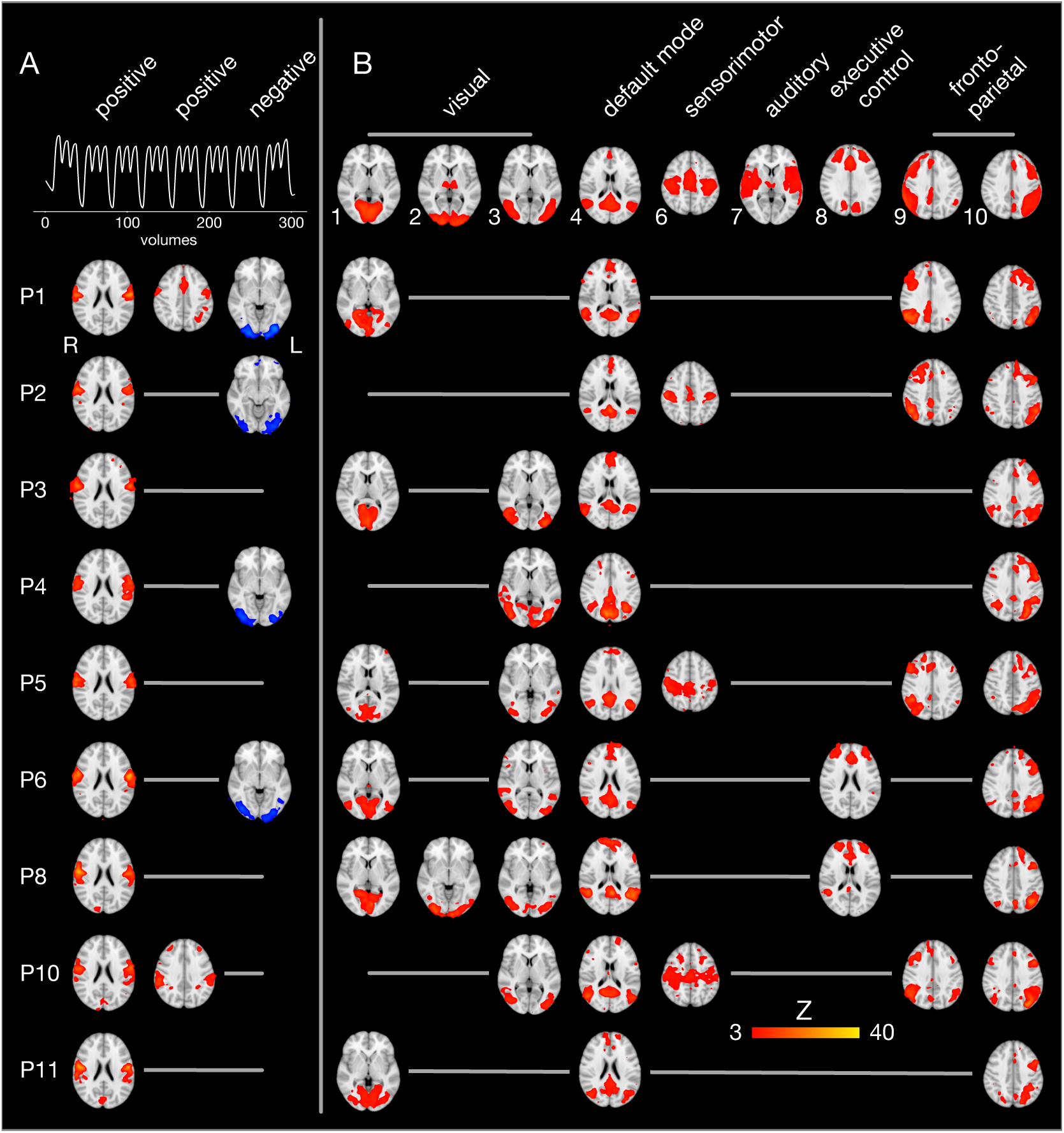
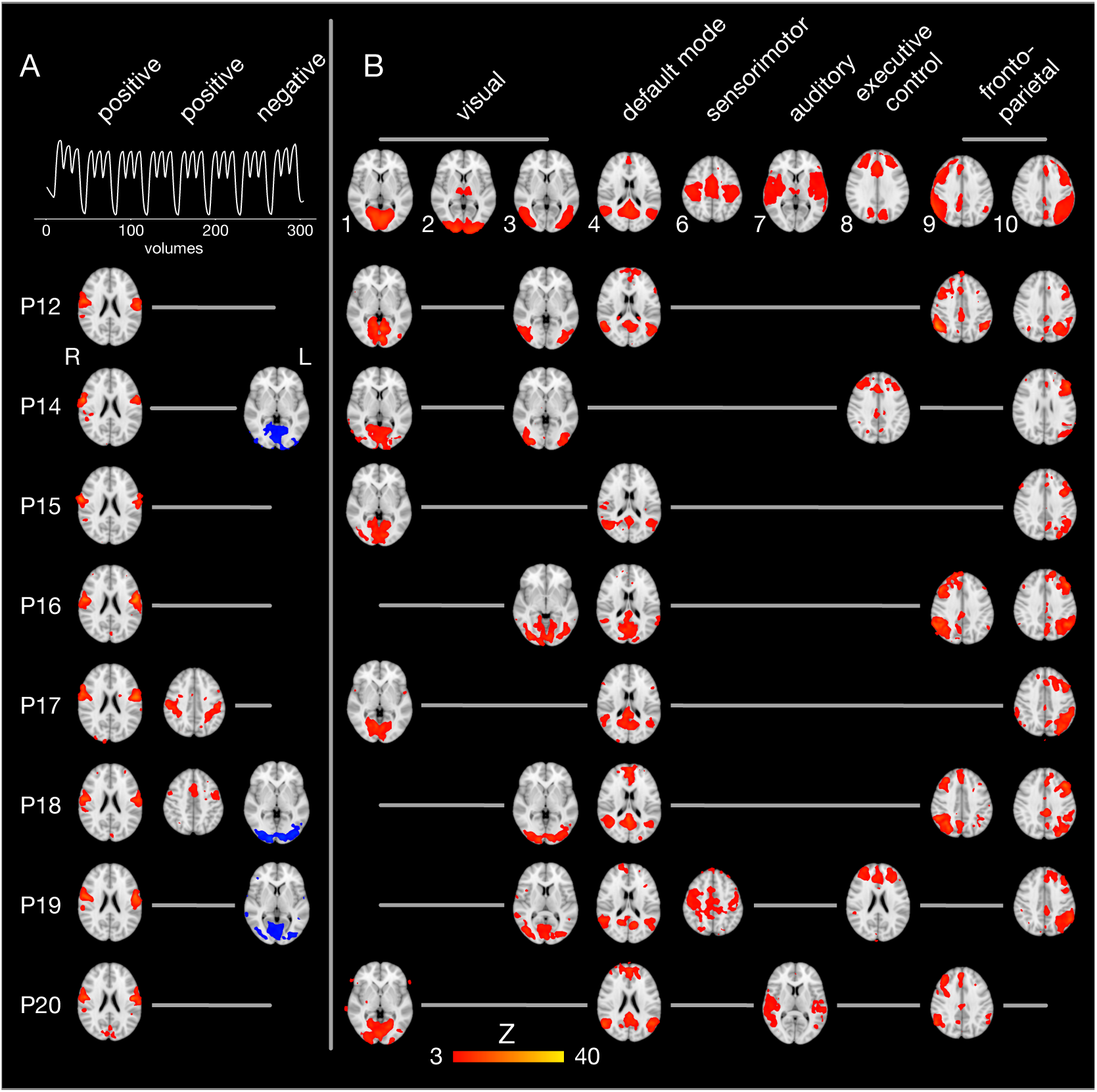
Spatio-temporal filtering of individual ICA components. The results of the single-session independent component analyses of all 17 participants (one participant per row) are summarized. Brain images are shown in radiological convention (the left hemisphere is seen on the right) after registration to the MNI152 standard space template. (**A**) The three columns on the left display components whose time course is significantly correlated with the expected brain activity (first row). The first and second column show positively correlated (r > 0.4, p < 0.05), the third column negatively correlated components (blue, r < −0.4, p < 0.05). (**B**) The upper row illustrates nine of ten established brain networks identified in resting state fMRI data (Smith et al., 2009). For every single participant, independent components are displayed that are spatially correlated with one of the established networks. Network 5 from the study by Smith et al. (2009) (consisting of large parts of the cerebellum) is not shown because it was not significantly correlated with cerebellar activity in one of the present fMRI data sets.

Spatial filtering of independent components revealed inter-individually variable patterns of brain activity during tongue movements. Spatial cross-correlations with established neural networks (shown in the first row of **Figure 4B**) (Smith et al., 2009) identified between one and three visual networks in all participants (except P2, who showed a visual component after temporal filtering). In addition, the default mode network and the left fronto-parietal network were active in all but one participant. Of note, the default mode network was not negatively correlated with the expected brain activity associated with tongue movements. The right fronto-parietal network was active in eight participants and the executive control network in four participants. In four participants, a superior sensorimotor network, comprising the hand sensorimotor cortex and the supplementary motor area, was found.

## 4 DISCUSSION

Main results of the present task-based fMRI study on the neural correlates of tongue movements were: (1) All three tongue movements under investigation were controlled by the same neurofunctional system, consisting of the bilateral tongue primary sensorimotor cortex, supplementary motor cortex, anterior cingulate gyrus, basal ganglia, thalamus, and cerebellum. (2) Distinct tongue movements also involved more specialized regions, such as the prefrontal, posterior parietal, and temporal cortices. (3) Using a novel approach to characterize inter-individual differences in task-based fMRI data, *spatio-temporal filtering of independent components*, we found consistent activation of the tongue primary sensorimotor cortex in all participants, but also remarkable variability, e.g. in fronto-parietal and executive control networks.

### 4.1 Model-based group fMRI analysis

The present study demonstrates the core cortical (lateral primary motor cortex (Fesl et al., 2003), supplementary motor area) and subcortical regions (basal ganglia, thalamus, cerebellum) of the tongue motor system, corroborating several previous fMRI studies (Brown et al., 2008; Corfield et al., 1999; Malandraki et al., 2009; Martin et al., 2004; Shinagawa et al., 2003; Watanabe et al., 2004). The tongue motor system was very similar for all three tongue movements under investigation (**Figure 3**). Our results also demonstrate the involvement of the lateral primary somatosensory cortex, reflecting the extensive mechanosensory (Kaas et al., 2006) and proprioceptive innervation of the tongue (Adatia and Gehring, 1971).

Moreover, the bilateral insular cortex was active during all three tongue movements. The insulae are not regarded as motor areas *per se*, but as areas of polymodal sensory, motor, cognitive, and affective integration. The insular cortex is involved in processing somatosensory (Pugnaghi et al., 2011; Sörös et al., 2008), gustatory (Small, 2010), and nociceptive stimuli (Xu et al., 2019). In addition, insular activity is associated with voluntary and semi-voluntary oro-facial movements, such as jaw opening and closing (Wong et al., 2011), speech production (Simonyan and Fuertinger, 2015; Tourville et al., 2019), and swallowing (Leopold and Daniels, 2010; Malandraki et al., 2011; Sörös et al., 2009). Importantly, insular activity is not specific for oro-facial movements, but has been found in simple finger movements as well (Turesky et al., 2016).

Tongue motor control was also associated with activity in prefrontal areas, critical for motor planning (Svoboda and Li, 2018), and in posterior parietal areas, involved in processing and perception of action-related information (Culham and Valyear, 2006). Parietal activity has also been found in previous fMRI studies during frontal tongue movements (tapping of the tip of the tongue against the alveolar ridge) (Malandraki et al., 2009) and a series of spatially complex tongue movements (pressing the inside of a left or right, upper or lower incisor, canine, or molar tooth with the tip of the tongue) (Watanabe et al., 2004).

Comparing neural activity between different tongue movements resulted in complex patterns of activation differences (**Table 3**). Speech-related frontal movements were associated with less activation in parts of the bilateral superior parietal lobule (vs. horizontal movements) and in parts of the bilateral precentral gyrus, right postcentral gyrus, and the left posterior parietal cortex (vs. vertical movements). We may speculate that, in most humans, speech-related tongue movements are probably the most overlearned movements and therefore are performed with less neural resources than less extensively trained tongue movements. Remarkably, frontal tongue movements were associated with increased activity in parts of the left frontal and temporal lobes compared with vertical movements. Again, we may speculate that frontal tongue movements, usually performed in the context of overt speech production, activate areas critical to speech-language production, such as the left inferior frontal gyrus, even when performed in isolation.

### 4.2 Model-free individual fMRI analysis

Almost all task-based and resting-state fMRI studies only present group analyses of neural activity. Recently, individual differences in brain structure and function have attracted growing attention (Dubois and Adolphs, 2016; Finn et al., 2015; Kanai and Rees, 2011). The study of individual differences of the neural control of tongue movements is expected to improve our understanding of tongue motor impairment, e.g. in amyotrophic lateral sclerosis (Kollewe et al., 2011) or Parkinson’s disease (Van Lieshout et al., 2011), and help in the assessment of the efficacy of treatment options such as tongue motor training (Arima et al., 2011; Komoda et al., 2015).

To investigate inter-individual differences in tongue movement-related neural activity, we used a novel approach, *spatio-temporal filtering of independent components*. First, ICA was employed to perform a low-dimensional decomposition of every single fMRI data set of 9:21 min duration. ICA carried out a model-independent separation of the original data into components that are related to neural activity, physiological extra-cerebral processes (such as respiration or blood pulsation), and imaging artifacts (such as head motion or susceptibility artifacts). The primary advantage of a model-free ICA was the detection of previously unexpected patterns of neural activity (McKeown et al., 1998). In the traditional general linear model-based analysis of fMRI time series, these patterns of activity would be considered as noise and discarded (Monti, 2011).

Second, we used a temporal filter to identify independent components whose time course was correlated with the expected brain response. This approach demonstrated that the activation of the lateral primary sensorimotor cortex was found in all 17 participants. Tongue movement is, similar to e.g. finger tapping, a very robust sensorimotor paradigm.

Third, we applied a spatial filter, i.e. we cross-correlated the independent components obtained for our participants with a set of ten established components (interpreted as neural networks). These canonical networks were derived from resting state fMRI data (Smith et al., 2009). Interestingly, nine of ten canonical networks were also found in our task-based data sets. In 16 participants, the default mode network was active, a large-scale neural network, including the posterior cingulate cortex, precuneus, medial prefrontal gyrus, and inferior parietal cortex (Greicius et al., 2003). In 16 participants, the left fronto-parietal network and in 8 participants, the right fronto-parietal network was active. Fronto-parietal networks have been shown to subserve attentional mechanisms (Markett et al., 2014) and are probably involved in a wide variety of tasks. In conclusion, *spatio-temporal filtering of independent components* appears to be a powerful tool to study inter-individual differences in brain function. Future studies are warranted to correlate the individual activation of neural networks with individual behavioral, clinical, and genetic information.

## AUTHOR CONTRIBUTIONS

PS and SS designed the study, acquired MRI data, and performed the data analysis. SS recruited the participants. PS prepared the figures and wrote the manuscript. SS and KW revised the manuscript and participated in the interpretations of the findings. PS conceived the *spatio-temporal filtering of independent components* approach.

## ACKNOWLEDGMENTS

The authors wish to thank Katharina Grote and Gülsen Yanc for assisting with MRI data acquisition.

## Footnotes

1 http://www.vislab.ucl.ac.uk/cogent_graphics.php

2 https://uol.de/en/medicine/biomedicum/neuroimaging-unit

3 http://stnava.github.io/ANTs/

4 https://www.oasis-brains.org

5 https://fsl.fmrib.ox.ac.uk/fsl/fslwiki/

6 https://github.com/maartenmennes/ICA-AROMA

7 https://fsl.fmrib.ox.ac.uk/fsl/fslwiki/FNIRT

8 https://fsl.fmrib.ox.ac.uk/fsl/fslwiki/Atlases

9 https://www.fmrib.ox.ac.uk/datasets/brainmap+rsns/

